# Genetic structure of prey populations underlies the geographic mosaic of arms race coevolution

**DOI:** 10.1101/585851

**Authors:** Michael T.J. Hague, Amber N. Stokes, Chris R. Feldman, Edmund D. Brodie, Edmund D. Brodie

## Abstract

Reciprocal adaptation is the hallmark of arms race coevolution, but the symmetry of evolutionary change between interacting species is often untested, even in the best-studied battles of natural enemies. We tested whether prey and predator exhibit symmetrical local co-adaptation in the example of a geographic mosaic of coevolution between toxic newts (*Taricha granulosa*) and resistant garter snakes (*Thamnophis sirtalis*). Prior work showing a tight correlation between levels of newt toxin and snake resistance is regarded as textbook evidence of the intense arms race between natural enemies. Here, we similarly found that toxin and resistance are functionally matched in prey and predator populations, further suggesting that mosaic variation in the armaments of both species results from the local pressures of reciprocal selection. Contrary to conventional wisdom, however, we found that local variation in newt toxin is best predicted by neutral population divergence rather than the resistance of co-occurring predators. Snake resistance, on the other hand, is clearly explained by local levels of prey toxin. Prey populations seem to structure variation in defensive toxin levels across the geographic mosaic, which in turn determines selection on predator resistance. Exaggerated armaments suggest that coevolution occurs in certain hotspots, but our results imply that neutral processes like gene flow—rather than reciprocal adaptation—structure the greatest source of variation across the landscape. This pattern supports the predicted role of “trait remixing” in the geographic mosaic of coevolution, the process by which non-adaptive forces dictate spatial variation in the interactions among species.

**SIGNIFICANCE STATEMENT:** When the weapons of natural enemies like prey toxins and predator resistance are matched across the geographic landscape, they are usually presumed to result from arms race coevolution. In the textbook example of an arms race, matched levels of newt toxin and garter snake resistance have long been regarded as evidence of such local co-adaptation. To the contrary, we found that local variation in newt toxicity is best explained by the neutral geographic structure of newt populations. This spatial variation of prey in turn dictates local selection on garter snakes, structuring the geographic pattern of predator resistance. These results demonstrate how landscape patterns of phenotypic variation are determined by a mixture of natural selection, historical biogeography, and gene flow that comprise the geographic mosaic of coevolution.

## INTRODUCTION

Coevolutionary dynamics result from the reciprocal selection generated by ecological interactions between species, which drives adaptation and counter-adaptation in the traits mediating interactions (1–4). Because the nature of species interactions and their fitness consequences vary spatially, heterogeneity in the form of reciprocal selection is expected to generate a geographic mosaic of coevolution, such that traits at the phenotypic interface of coevolution covary due to local conditions (2, 5–7). Coevolving species are expected to exhibit matched trait variation across the landscape, for example, in seed traits and their predators (5, 7) or host and pathogen genotypes (8, 9). As such, local co-adaptation is often presumed when a pair of interacting species share matched abilities throughout their shared range (1, 2, 4).

However, a geographic mosaic of matched phenotypes may not result solely from reciprocal co-adaptation (10, 11). A simpler, non-adaptive explanation for matched trait variation involves the spatial population structure and ancestry of each species. Common barriers to dispersal or a shared biogeographic history, for example, could structure phenotypic divergence congruently in co-occurring species (12). Populations are subject to selectively neutral processes, such as drift and gene flow, that continually alter the spatial distribution of alleles and traits at the phenotypic interface, a process termed “trait remixing” in the geographic mosaic theory of coevolution (2, 10, 13). Only when trait variation deviates from the neutral expectations of population structure in both species can we infer local co-adaptation in the geographic mosaic of coevolution (10). Otherwise, phylogeography, drift, and gene flow provide a parsimonious explanation for patterns of divergence across the landscape.

In the geographic mosaic of arms coevolution, populations of rough-skinned newts (*Taricha granulosa*) and common garter snakes (*Thamnophis sirtalis*) tend to have matched levels of prey toxin and predator resistance, a pattern generally interpreted as evidence of local co-adaptation (11, 14–16). We tested whether this inferred signature of coevolution can instead be explained by the population structure of either species. In western North America, rough-skinned newts secrete the deadly neurotoxin tetrodotoxin (TTX), which binds to the outer pore of voltage-gated sodium channels (Na_V_) and prevents the initiation of action potentials (17, 18). Common garter snakes exhibit resistance to TTX that is largely due to specific amino acid substitutions in the fourth domain pore-loop (DIV p-loop) of the skeletal muscle sodium channel (Na_V_1.4) that disrupt toxin-binding (Fig. 1) (19, 20). Channel-level TTX resistance conferred by each allele in the DIV p-loop is tightly correlated with muscle and whole-animal levels of phenotypic resistance (19–21). TTX-resistant alleles occur at high frequency in coevolutionary “hotspots” with highly toxic newts, but are largely absent in surrounding “coldspots” where newts are non-toxic (22). Because quantitative levels of newt toxin and snake resistance are functionally matched in most populations, the geographic mosaic of coevolution is thought to primarily consist of co-adaptations to local pressures in the arms race (11, 14). If true, we predict that population divergence in both prey toxin and predator resistance represent adaptive deviations from the selectively neutral expectations of population structure and ancestry.

**Fig. 1.**
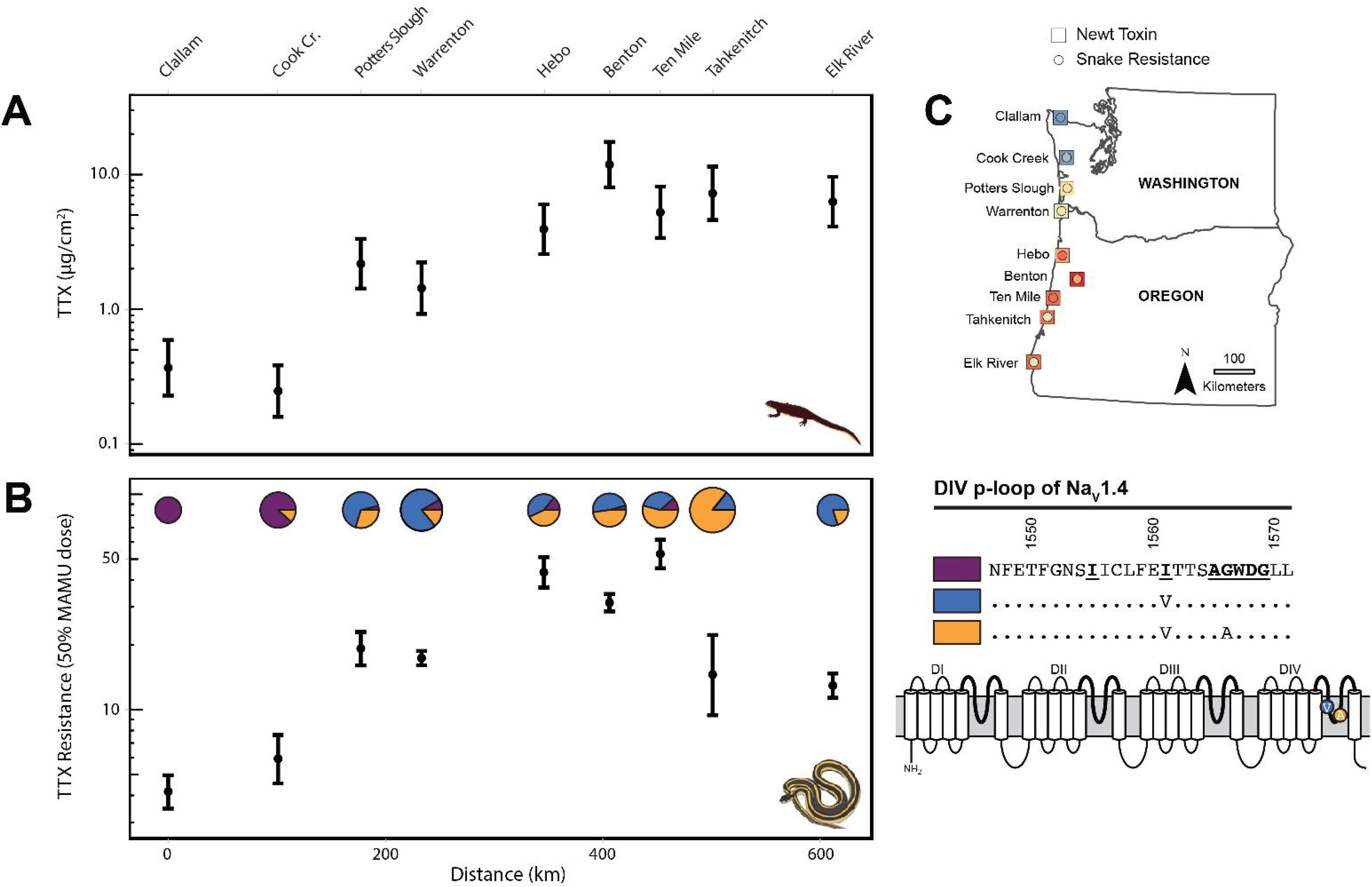
Matching phenotypes in prey and predator imply local co-adaptation in the arms race. (**A**) Population means of TTX levels (μg/cm^2^) in newts and (**B**) phenotypic TTX resistance (50% MAMU dose) in snakes along the latitudinal transect. Error bars indicate 95% confidence intervals. The x-axis represents linear distance (km) from the northernmost sampling site (Clallam; 0 km). For snakes, the frequency of TTX-resistant alleles in the Na_V_1.4 channel is shown with pie charts proportional to sample size. To the right, the schematic of Na_V_1.4 shows the four domains of the channel (DI–DIV), with the extracellular pore loops (p-loops) highlighted with bold lines. Specific amino acid changes in the DIV p-loop are shown in their relative positions within the pore. The TTX-sensitive ancestral sequence (purple) is listed, followed by the two derived alleles known to confer increases in channel resistance in this lineage. (**C**) Map inset illustrates population estimates of prey toxin and predator resistance at each location in the geographic mosaic. Blue colors correspond to low estimates of TTX (squares) or resistance (circles), whereas red indicates escalated phenotypes in the arms race.

We conducted fine-scale population sampling of newts (*n*=138) and garter snakes (*n*=169) along a latitudinal transect of nine locations on the Pacific Coast in Washington and Oregon (USA) that spans the geographic mosaic (Fig. 1, Supplemental Table S1), ranging from low ancestral levels of newt toxin and snake resistance (northern Washington) to a hotspot of extreme escalation in both species (central Oregon). We focused our sampling in Washington and Oregon, because other regions of sympatry, particularly California, contain additional TTX-bearing newt species (e.g., *Ta. torosa*) and resistant garter snakes (e.g., *Th. atratus*) that confound inferences of coevolution between discrete pairs of species. At each sampling site in the mosaic, we characterized levels of TTX in newt populations and TTX resistance in garter snakes, including whole-animal phenotypic resistance and Na_V_1.4 channel genotypes. We then tested whether trait divergence in the weaponry of each species deviates from the neutral expectations of geographic population structure using genome-wide single nucleotide polymorphisms (SNPs). If the geographic mosaic of prey toxin and predator resistance results from co-adaptation to local pressures in the arms race, then trait divergence for each species should be predicted solely by the local armaments of its natural enemy. Alternatively, if mosaic variation in toxin and/or resistance is a selectively neutral result of population structure, then trait divergence should be better predicted by genomic SNP variation (23, 24).

## RESULTS AND DISCUSSION

### Matched trait variation implies local co-adaptation

Geographic patterns of newt TTX and snake resistance were broadly consistent with previous work indicating arms race coevolution has led to the closely matched phenotypes in each species (11, 14). TTX levels (μg/cm^2^) of newts varied by population (ANOVA; F[8,114]=37.43, p<0.001) and by sex (F[1,114]=4.37, p=0.039) along the latitudinal transect (Fig. 1; Supplemental Table S1). TTX resistance (50% MAMU dose) of snakes also varied by population (according to non-overlapping 95% confidence intervals; Fig. 1; Supplemental Table S1) and was closely correlated with prey toxin (regression of distance matrices, p=0.021; Table 1). The presence of TTX-resistant alleles in the Na_V_1.4 channel co-varied with phenotypic resistance in garter snakes, such that pairwise F_ST_ divergence at the DIV p-loop was correlated with population divergence in phenotypic resistance (Mantel test, r=0.47, p=0.032).

**Table 1.** Results from multiple regression of distance matrices (MRMs) comparing population divergence in phenotypic and genetic data.

| Response Variable | Explanatory Variable(s) | Coefficient | p-value | R<sup>2</sup> |
| --- | --- | --- | --- | --- |
| Newt TTX levels | Neutral F<sub>ST</sub> of newts | 5.998 | **0.002\*** | 0.414 |
|  | TTX resistance of snakes | 0.415 | **0.019\*** | 0.274 |
|  | Neutral F<sub>ST</sub> + TTX resistance |  |  |  |
|  | Neutral F<sub>ST</sub> of newts | 4.827 | **0.006\*** | 0.501 |
|  | TTX resistance of snakes | 0.253 | 0.065 |  |
| Snake TTX resistance | Neutral F<sub>ST</sub> of snakes | 1.442 | 0.719 | 0.006 |
|  | TTX level of newts | 0.662 | **0.021\*** | 0.274 |
|  | Neutral F<sub>ST</sub> + TTX toxicity |  |  |  |
|  | Neutral F<sub>ST</sub> of snakes | -5.830 | 0.189 | 0.338 |
|  | TTX level of newts | 0.873 | **0.011\*** |  |
|  | Neutral F<sub>ST</sub> of newts | 4.632 | **0.035\*** | 0.155 |
| Snake F<sub>ST</sub> of DIV p-loop | Neutral F<sub>ST</sub> of snakes | 2.649 | 0.202 | 0.080 |
|  | Neutral F<sub>ST</sub> of newts | 3.196 | **0.010\*** | 0.311 |

Quantitative estimates of prey toxin and predator resistance at each locality showed functional overlap in the armaments of each species, implying that locally varying reciprocal selection occurs throughout the geographic mosaic (Fig. 2) (11). For each locality, we estimated whole-newt levels of TTX (mg) and the dose of TTX (mg) required to reduce the performance of co-occurring snakes by 15%, 50%, and 85% (see Methods). The TTX dose required to reduce snake performance by 50% is considered a perfect functional match between newt TTX and snake resistance. The 15% and 85% doses delimit the range of functional overlap between toxin and resistance, outside of which prey and predator are considered so mismatched that variable fitness outcomes and reciprocal selection are unlikely to occur (11, 15, 16). Under this framework, each locality we sampled exhibited some degree of overlap in the phenotypic distributions of newt TTX and snake resistance, indicating potential for reciprocal selection between prey and predator. Interestingly, all point estimate comparisons of TTX and resistance fell below the 50% line, suggesting that garter snakes on average tend to have higher levels of resistance than the toxins of co-occurring newts, a pattern consistent with the prediction that predators experience intense selection when prey are deadly (see below) (3). Nonetheless, under this framework of previous work, our results imply that matched trait variation in the geographic mosaic of coevolution may be a result of reciprocal selection. Next, we characterized neutral genomic variation for each species to test whether population genetic structure can alternatively explain this observed pattern of matched weaponry.

**Fig. 2.**
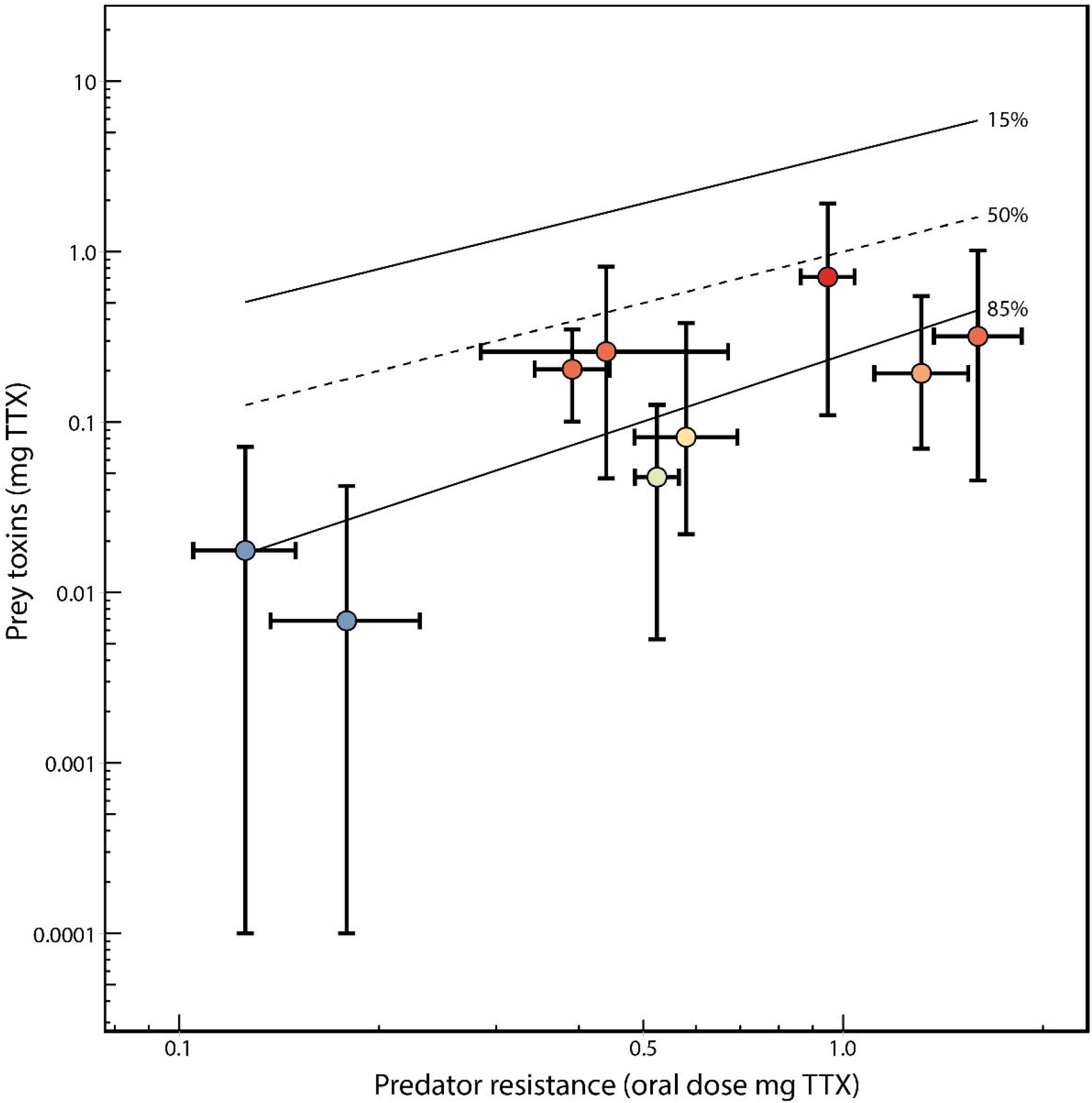
Functional overlap in levels of prey toxin and predator resistance. The functional relationship of average adult newt total skin TTX (mg, log scale) is plotted against the oral dose of TTX (mg, log scale) required to reduce the speed of an average adult garter snake to 50% of baseline speed post-ingestion of TTX for each locality. Individual points are colorized by the average level of newt TTX as in Fig. 1C. Vertical bars represent the full range of newt TTX values observed for each population; horizontal bars represent the 95% confidence interval for the oral 50% dose of the corresponding snake population. The dashed 50% line represents a functional match between newt TTX and snake resistance: the dose of TTX in a newt that would reduce a snake of a given resistance to 50% of its performance. The solid 15% and 85% lines (calculated as best-fit regressions for each locality) delimit the range of functionally relevant TTX doses for snakes across the range of sampled localities. Below the 85% line, values of TTX resistance are high enough for snakes to consume co-occurring newts with no reduction in performance or fitness. Above the 15% line, doses of newt TTX are so high that any snake that ingests a newt would be completely incapacitated or killed. Localities with error bars that fall *entirely* outside the boundaries of these lines (none of which were observed) are considered mismatched and regions where reciprocal selection could not occur.

### Prey and predator populations differ in geographic structure

We used neutral SNPs generated from double digest restriction-site associated DNA sequenced (ddRADseq; see Methods) to characterize the population structure of prey and predator, which served as a neutral expectation for trait divergence in the geographic mosaic of coevolution. Global F_ST_ values for newts (F_ST_=0.068, 95% CI [0.065, 0.069]) and snakes (F_ST_=0.070, 95% CI [0.067, 0.075]) were similar, and both species exhibited a pattern of isolation-by-distance (IBD) along the transect (distance-based redundancy analysis; newts, F=-38.528, p=0.002; snakes, F=22.021, p=0.001). This pattern is consistent with previous work suggesting limited population structure and a recent northward post-glacial expansion of both species (25–27). Pairwise estimates of F_ST_, however, revealed subtle differences in the geographic population structure of prey and predator (Supplementary Table S4), a pattern that was also supported by principal coordinate (PCoA) and Bayesian clustering (STRUCTURE) analyses (Fig. 3, Supplemental Fig. S1). In particular, the major axis of variation from the PCoA (PCo1) and STRUCTURE revealed limited structure among newt populations along the transect, whereas garter snakes showed evidence of population subdivision near the Washington-Oregon border. For example, the STRUCTURE plot for snakes indicates that subdivision between the two most likely genetic clusters (K=2) occurs just south of the Columbia River near the Warrenton locality in Oregon. Because pairwise F_ST_ estimates, the PCoAs, and STRUCTURE produced very similar results, yet rely on different assumptions, we feel confident in our characterization of population structure for each species. Finally, we tested whether these patterns of population structure provide a more parsimonious explanation than local co-adaptation for the geographic mosaic of matched trait variation between prey and predator.

**Fig. 3.**
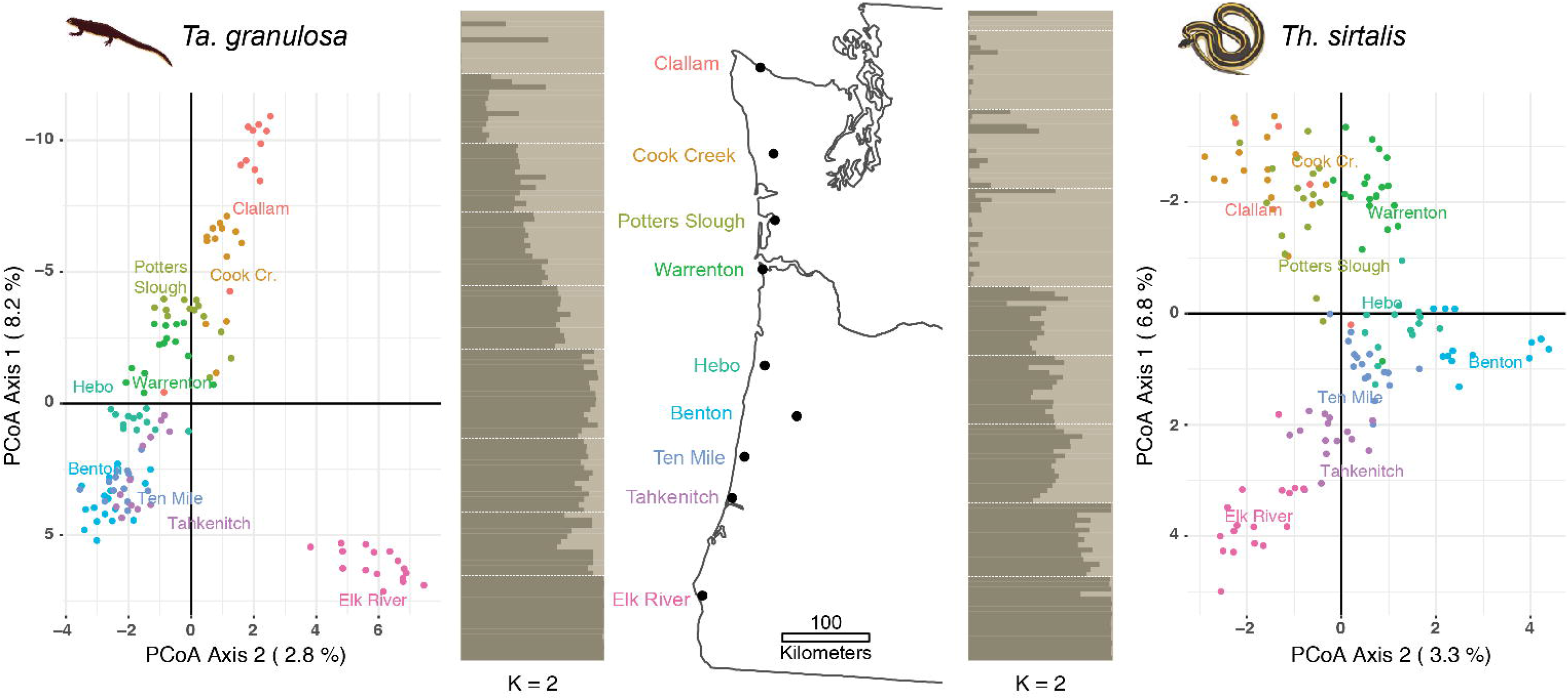
Populations of prey and predator differ in geographic structure. Results from the principal coordinate (PCoA) and STRUCTURE analyses of neutral SNPs from newts and snakes. PCoA graphs are rotated 90° to emphasize the major axis of variation corresponding to latitude. The PCo1 values for each individual were used as a neutral expectation in the cline-fitting analyses. STRUCTURE plots are arranged latitudinally by population, in the same order as the map. Each horizontal bar represents the ancestry assignment of an individual, with populations separated by white dashed lines.

### Asymmetric evidence of local adaptation in prey and predator

Trait divergence in newt TTX was best predicted by population genetic structure, not predator resistance, suggesting that reciprocal co-adaptation cannot fully explain the geographic mosaic of matched toxin and resistance. We generated distance matrices to test whether phenotypic divergence in one species (e.g., newt TTX levels) is best explained by (1) neutral genomic divergence (pairwise F_ST_; Supplemental Table S4) or (2) phenotypic divergence in the natural enemy (snake resistance). In univariate regressions, population divergence in the TTX level of newts was strongly predicted by neutral F_ST_ divergence (R^2^=0.414), as well as TTX resistance of garter snakes (R^2^=0.274; Table 1). F_ST_ divergence remained significant in the multiple regression, indicating that population structure of newts predicts TTX levels, even after controlling for TTX resistance of the predator (which was at best marginally significant; p=0.065). Unlike newt TTX, garter snake resistance was strictly predicted by the prey toxin and not neutral F_ST_ divergence. Both phenotypic resistance and F_ST_ divergence at the DIV p-loop (the site of toxin-binding in Na_V_1.4) were uncorrelated with neutral F_ST_ values in garter snakes (Table 1). These results indicate that neutral genetic divergence is a key determinant of population differences in prey toxin levels, which in turn predicts differences in TTX resistance among predator populations. The geographic structure of newt populations appears to be so influential to spatial dynamics that divergence in garter snake phenotypic resistance and F_ST_ at the DIV p-loop are both significantly predicted by neutral F_ST_ divergence of newts (Table 1).

Clinal variation in TTX levels of newts is highly congruent with neutral genomic structure based on the major axis of variation from the PCoA (PCo1; Fig. 4) and Bayesian clustering analyses (Supplemental Fig. S2). TTX resistance of garter snakes, in contrast, clearly deviates from neutral expectations to track variation in prey toxin. Cline-fitting analyses show that prey toxin levels and predator resistance are tightly matched along the 611 km transect; the geographic center points of each cline are located just 64 km apart and do not differ statistically. The cline center of TTX-resistant alleles in snakes is also located nearby, although it differed statistically from the center of newt TTX. Despite similar phenotypic clines for both species, variation in levels of newt toxin showed an even tighter match to clinal variation in neutral population structure. The center points of the TTX and neutral clines were located only 19 km apart. PCo1 from the PCoA was a strong predictor of variation in TTX levels (linear model; t-value=5.682, p<0.001), even after controlling for the effect of TTX resistance of garter snakes (Supplemental Table S6, model 3). Conversely, variation in phenotypic resistance and TTX-resistant alleles in snakes both deviated significantly from the neutral cline (Fig. 4), such that resistance was not predicted by PCo1 or 2 from the PCoA (Supplemental Table S6, model 3). The center points of the snake phenotypic resistance and neutral clines were located a distant 310 km apart.

**Fig. 4.**
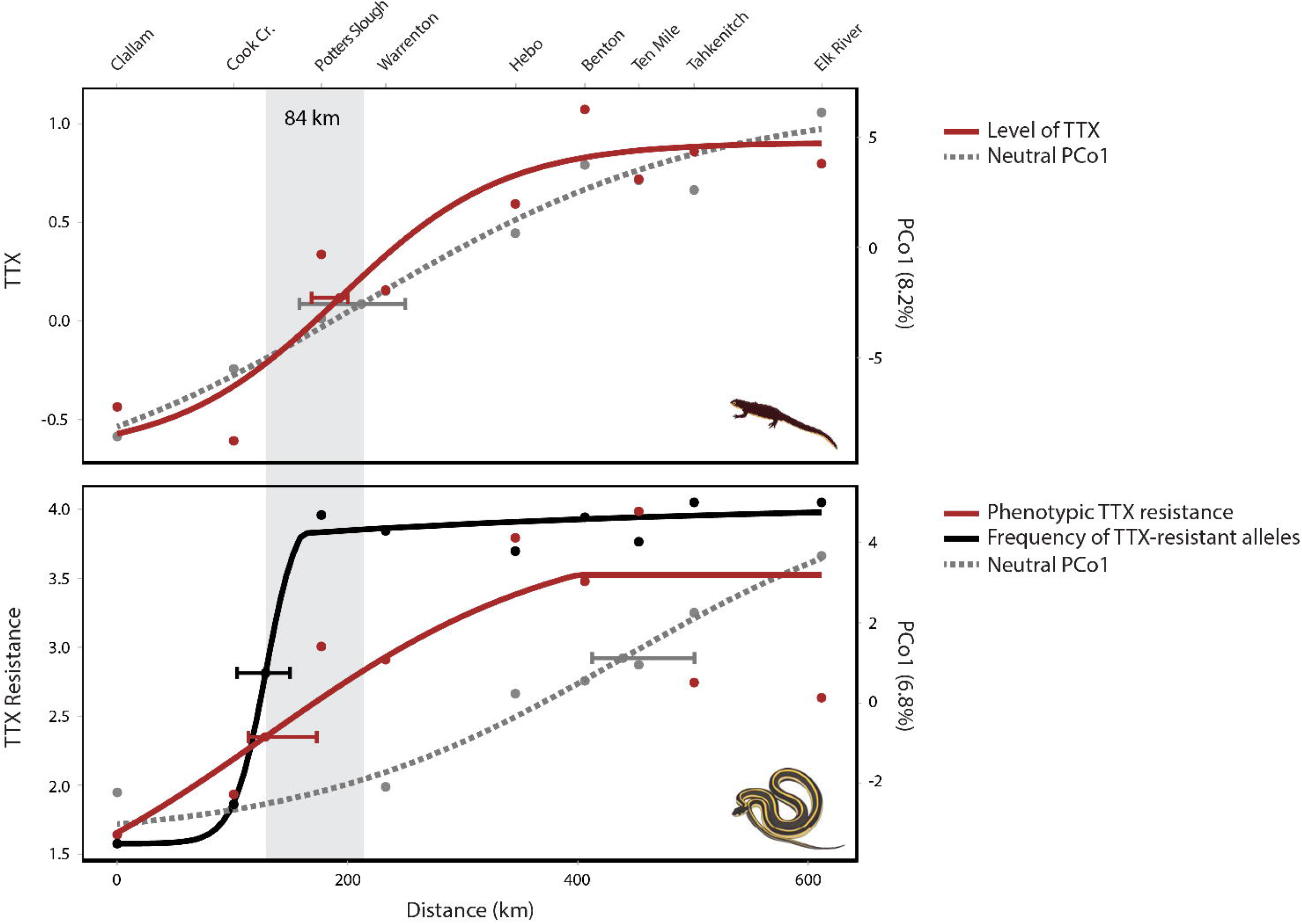
Levels of prey toxin are best predicted by neutral population structure, whereas predator resistance is predicted by prey toxin. Cline-fitting results for phenotypic and genetic variation are shown, with error bars indicating confidence intervals surrounding the geographic cline centers. (**A**) Phenotypic clines of TTX levels (log[TTX μg/cm^2^ + 0.1]) and (**B**) TTX resistance (ln[MAMU + 1]) are shown in red. For snakes, the frequency of TTX-resistant alleles in the Na_V_1.4 channel was also modeled (in black). Gray dashed lines represent the neutral expectation for trait variation due to population structure, based on the PCoA. The cline center points of TTX levels and neutral PCo1 in newts, and phenotypic resistance and TTX-resistant alleles in snakes, are all located within in 84 km of each other along the 611 km transect.

### Implications for the geographic mosaic of arms race coevolution

Levels of prey toxin and predator resistance are tightly matched across the landscape, but this pattern does not appear to primarily result from local arms race co-adaptation. Although predator resistance is geographically structured by a signature of local adaptation to prey, levels of the prey toxin are structured in tight accordance with neutral population divergence. These results imply that local variation in the newt toxin across the geographic mosaic is best explained by non-adaptive processes, such as drift and historical biogeography, rather than local variation in selection from predators. For example, latitudinal patterns of newt TTX and neutral divergence are both consistent with a signature of isolation-by-distance along the transect (25, 26).

The asymmetric signatures of adaptation we observed in prey and predator may reflect differences in the mechanisms that underlie phenotypic variation in each species. Genes associated with TTX biosynthesis have yet to be discovered; however, toxin production in newts likely requires a complicated biosynthetic pathway. For example, biosynthesis of a similar neurotoxin found in puffer fish, saxitoxin (STX), involves gene expression in a cluster of up to 26 genes (28). Some researchers suggest exogenous factors, like environmentally-derived precursors, may also affect the ability of newts to synthesize or sequester TTX (29, 30). On the other hand, TTX resistance in garter snakes is largely due to a small number of amino acid changes to the p-loops of the Na_V_1.4 channel (19–22). These large-effect mutations could make TTX resistance more evolutionarily labile than the defensive toxins in newts, permitting rapid local adaptation in predator populations (11, 21).

Asymmetric evolutionary changes could also arise from a selective imbalance associated with the interactions between prey and predator. In antagonistic interactions, the species under more intense selection is generally expected to be better adapted to local conditions (31). While prey are typically thought to experience stronger selection than their predators (the “life-dinner principle”) (32), this asymmetry may be reversed when prey contain deadly toxins like TTX (3). In fact, populations in central Oregon are the most toxic newts known (11), so non-resistant predators should experience severe fitness consequences. Moreover, we found that garter snakes tend to have greater functional estimates of resistance than the levels of TTX in co-occurring newts (Fig. 2), implying intense selection on predators.

Non-adaptive processes of drift and gene flow provide a parsimonious explanation for the overall pattern of variation in newt TTX across the geographic mosaic, but the extreme levels of prey toxin and predator resistance at phenotypic hotspots such as central Oregon are likely a result of arms race coevolution. After controlling for the effects of population structure, escalation of TTX in newts was still marginally predicted by the TTX resistance of local garter snakes in MRM analyses (p=0.065; Table 1) and significantly predicted by TTX resistance in a linear model (t-value=2.803, p=0.006; Supplemental Table S6, model 3). This relationship seems to be primarily driven by the southern portion of the transect, where intense reciprocal selection has presumably led to the escalation of armaments in both species. Taken together, the overall pattern suggests that co-adaptation can explain escalated TTX levels in specific hotspots of coevolution, but global variation in TTX across the geographic mosaic is dictated by processes like drift and gene flow. For example, TTX may be favored in specific hotspots of coevolution, but not in surrounding regions like northern Washington (i.e., “coldspots”), and the tight spatial correlation we observed between newt TTX and neutral SNPs reflects gene flow homogenizing variation between hot- and coldspots.

The underlying importance of prey population structure in the geographic mosaic of coevolution points to an influential role of “trait remixing”, a largely untested component of the geographic mosaic theory thought to generate spatial variation in the phenotypic interface of coevolution (2, 10). The neutral processes of drift and gene flow are predicted to continually alter the spatial distribution of allelic and phenotypic variation, potentially interfering with local selection. Gene flow outwards from hotspots of coevolution is predicted to alter dynamics in surrounding populations (13, 33), and if gene flow is high, the population with the strongest reciprocal effects on fitness is expected to dictate broader landscape patterns of trait variation (31, 33, 34). The homogenizing effects of gene flow may be less influential in snake populations due to the simple genetic basis of TTX resistance and/or strong selection on predators.

Our results highlight how not all landscape patterns of phenotypic matching in natural enemies may be the inherent result of coevolution (10). External factors such as abiotic conditions (35), evolutionary constraints (36), or interactions with other species (37) are likely to have unique effects on the evolution of prey and predator. In the geographic mosaic of arms race coevolution, it appears that phenotypic divergence in newt toxin in disproportionally the result of population structure, a pattern that underlies landscape-wide variation in the armaments of both species. Ultimately, the evolutionary response to selection at the phenotypic interface is almost certain to differ in two interacting species—so much so that coevolution may not always be the most parsimonious explanation for observed patterns of phenotypic divergence and trait matching across the geographic mosaic.

## ACKNOWLEDGEMENTS

We thank the Departments of Fish and Wildlife in Washington and Oregon for scientific collecting permits to MTJH (18-082 and 063-18, respectively). We also thank IACUC for the protocol to EDB, Jr. at USU (1008). We are grateful for T. St. Pierre’s help with fieldwork. J. McGlothlin and K. Gendreau helped with sex-linked analyses of Na_V_1.4 and provided valuable feedback that improved the quality of this manuscript. R. Cox, D. Taylor, A. Bergland, D. Carr, and the Brodie and Feldman lab groups also provided helpful input. This work was supported by a Doctoral Dissertation Improvement Grant from the National Science Foundation to MTJH and EDB III (DEB 1601296) and NSF support to EDB III (DEB 0922251).

## AUTHOR CONTRIBUTIONS

MTJH designed the project, collected specimens, generated genetic data, and performed statistical analyses. ANS collected phenotypic data on newt TTX levels. CRF collected specimens and phenotypic data on snake resistance. EDB, Jr. collected phenotypic data on snake resistance and provided leadership on the project. EDB III designed the project and provided leadership. All authors prepared the manuscript.

## METHODS

We sampled phenotypic and genomic data from *Ta. granulosa* (*n*=138) and *Th. sirtalis* (*n*=169) at nine locations along a latitudinal transect in the states of Washington and Oregon (Fig. 1; Supplemental Table S1) and compared mosaic patterns of escalation in the arms race to neutral population genomic structure across the landscape.

### TTX levels of Taricha granulosa

We estimated levels of TTX in *Ta. granulosa* using a Competitive Inhibition Enzymatic Immunoassay (CIEIA) and TTX-specific antibodies (38, 39). We quantified the amount of TTX in a 5mm circular skin punch from the dorsum of each newt using a human skin-biopsy punch (Acu-Punche, Acuderm Inc.) (26, 40, 41). These data were used to estimate the dorsal skin concentration of TTX (μg/cm^2^) in each individual. TTX is uniformly distributed throughout the dorsum and levels of TTX in the dorsal skin are tightly correlated with toxicity in other regions (41). We conducted a two-way ANOVA to test whether TTX differed by population and by sex, because past work suggests toxin levels may vary by sex (40). Distribution and leverage analyses indicated that a log(x + 0.1) transformation of TTX was needed. Transformed mean and variance values were used in the cline-fitting analysis of TTX along the transect.

Although a considerable body of work has described the genetic basis of TTX resistance in *Th. sirtalis* (see below), similar information regarding TTX in *Ta. granulosa* is unavailable. Although we were unable to characterize genetic variation underlying prey toxin, we assume that TTX production probably has a polygenic basis. For example, biosynthesis of saxitoxin (STX), a similar neurotoxin found in marine species, involves gene expression in a cluster of up to 26 genes in cyanobacteria (42).

### TTX resistance of Thamnophis sirtalis

We measured phenotypic TTX resistance using a well-established bioassay of whole animal performance (14, 22, 43, 44). Briefly, each individual was assayed on a 4 m racetrack to characterize its “baseline” crawl speed, then injected intraperitoneally with a known dose of TTX and assayed for “post-injection” speed. Population estimates of phenotypic TTX resistance are reported on a scale of mass-adjusted mouse units (MAMUs) to control for differences in body size (14). Resistance was estimated as the relative performance after injection: the MAMU dose of TTX that reduces performance by 50% of baseline speed. We incorporated racetrack data from previously published estimates of resistance from the same sampling locations to generate precise population estimates of phenotypic resistance in this study (see Supplemental Table S1) (14, 45).

The 50% MAMU dose was estimated separately for each population from a dose-response curve using curvilinear regression and the general transform 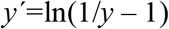 (14). Individuals from each population received a series of TTX doses, with an average of 2.5 different doses per individual. At *y*=0.5 (i.e., 50%), 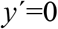 and the 50% dose is estimable 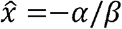 (where *α* is the intercept and *β* the slope from the curvilinear regression). Because 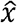 takes the form of a ratio, the standard error for the estimated 50% dose is calculated using standard methods for the variance of a ratio (14, 46). Confidence intervals of 95% were calculated as ±1.96 SE. Regression was performed in R with the “lmer” function implemented in the lme4 package (47). The individual ID of each snake was included as a random effect to account for the fact that each snake received multiple injections. Distribution and leverage analysis indicated that a transformation of the *x* variable (MAMU of TTX) was needed, so we transformed the data using *x*’=ln(*x* + 1) (14). Differences among populations in phenotypic TTX resistance were deemed significant if 95% confidence intervals did not overlap by more than half of a one-sided error bar (48).

We genotyped snakes for their amino acid sequence in the DIV p-loop of the Na_V_1.4 channel. Methods for Sanger sequencing are described in Hague et al. (2017) (22). For each individual, we sequenced a 666 bp fragment that includes the DIV p-loop region of Na_V_1.4. The Na_V_1.4 protein is encoded by the *SCN4A* gene located on the Z sex chromosome of *Th. sirtalis* (Gendreau et al., *in prep*). Colubrid snakes, including garter snakes, have heteromorphic sex chromosomes (ZZ males, ZW females) that are non-recombining (49, 50), and females appear to be hemizygous for the Z-linked *SCN4A* gene. We used sex-specific PCR primers to confirm the sex of each individual (ZW.734.F, 5’-TAAGGTCCTGGGCATGTCCT-3’; ZW.2468.R, 5’-ATGGCTTGGAATGAGGTGGG-3’; Gendreau et al., *in prep*). In DIV p-loop sequences from males, heterozygous positions on chromatograms were identified by eye and confirmed in both directions with sequencing. The haplotype phase of the DIV p-loop sequence for each male was inferred computationally with the program PHASE (51).

We translated the aligned DIV p-loop coding sequences into the amino acids and tested for departures from Hardy-Weinberg Equilibrium (HWE) using a joint test for HWE and equality of allele frequencies (EAF) using the *HWTriExact* function in the HardyWeinberg package in R (52–54). Standard tests for HWE rely on the assumption of EAF in males and females. The joint exact test for HWE and EAF accounts for the hemizygous sex and tests for joint departures of HWE and EAF (53–55). We also calculated pairwise F_ST_ divergence at the DIV p-loop in the program Arlequin (56) and used a Mantel test to test for a relationship between pairwise F_ST_ divergence at the DIV p-loop and phenotypic divergence in whole-animal TTX resistance. Significance was tested with 1000 permutations.

### Functional analysis of trait matching

Following Hanifin et al. (2008) (11), we estimated functional levels of newt TTX and snake resistance to visualize whether prey and predator exhibit matched levels of escalation along the transect. The model provides a rough estimate of functional interactions between newt TTX and snake resistance based on an extensive body of work (4, 14, 41, 43, 44, 57–59). Localities are considered “matched” if a sympatric interaction between prey and predator could potentially result in variable fitness outcomes for both species, leading to reciprocal selection between newt TTX and snake resistance (11). For each locality, we estimated whole-newt levels of TTX (mg) and the dose of TTX (mg) required to reduce performance of co-occurring snakes to 15%, 50%, and 85% of their baseline performance. The TTX dose required to reduce snake performance by 50% is considered a perfect functional match between newt TTX and snake resistance. The 15% and 85% doses delimit the range of functionally relevant doses for snakes. At performance levels <15%, all snakes that ingest newts are fully immobilized or killed and newts escape, whereas at performance levels >85% all snakes are unaffected and captured newts die (11, 15, 16). Localities where the full range of TTX doses found in newts fall outside the 15-85% region of phenotypic space are considered phenotypic “mismatches”, such that variable fitness outcomes and reciprocal selection are unlikely to occur.

Methods for estimating functionally comparable values of newt TTX and snake resistance are described in Hanifin et al. (2008) (11). Briefly, we extrapolated our measures of newt TTX in skin samples (μg/cm^2^) to the whole animal (mg of TTX/newt) using standard methods. For snakes, the 15, 50, and 85% MAMU doses were extracted from the dose response curves of each population (as described above). We converted snake resistance from MAMUs based on intraperitoneal (IP) injections to mg of TTX in oral doses using the following equation:

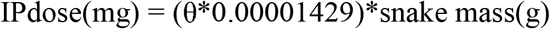

where θ is the MAMU dose and 0.0001429 is the conversion factor (1 MAMU= 0.01429μg TTX per gram of snake). The effects of TTX are dependent on body size, so we estimated the oral dose of TTX required to slow the average adult *Th. sirtalis* (mean adult mass = 52.84 g). We then converted the IP dose (mg) to the oral dose required to achieve the same performance reduction by multiplying the IP dose by 40. Estimates of orally ingested doses of TTX (mg) required to reduce snake performance by 15, 50, and 85% were used as a functionally equivalent metric to compare predator phenotypes to those of the prey (Supplemental Table S2).

### ddRADseq library preparation

We prepared ddRADseq libraries separately for *Ta. granulosa* and *Th. sirtalis* using the protocol described in Peterson et al. (2012) (60). Genomic DNA was extracted using the DNeasy Blood & Tissue kit (Qiagen Inc., Valencia, CA.). We digested 600 ng of genomic DNA for each sample using the restriction enzymes *EcoRI* and *SbfI* for *Ta. granulosa* and *MfeI* and *SbfI* for *Th. sirtalis*. Unique combinations of individual P1 and P2 barcoded adapters were annealed to the digested genomic DNA of each sample. Each barcode was six base pairs long and differed by at least two nucleotides. After barcoding, *Ta. granulosa* and *Th. sirtalis* samples were pooled separately, purified with AmpureXP beads (Beckman Coulter, Inc., Brea, CA, USA), and size selected for 500 to 600 bp fragments using a Pippin Prep (Sage Science, Inc., Beverly, MA, USA). We enriched the adapter-ligated fragments in the size-selected libraries using 16 PCR cycles and then purified the product with AmpureXP beads. The *Ta. granulosa* and *Th. sirtalis* libraries were each run on two lanes of the Illumina HiSeq 2500 platform (Illumina, Inc., San Diego, CA, USA) to generate 125 bp paired-end reads.

### Data processing of Illumina sequencing

Read quality of the raw sequence data was assessed using FastQC 0.11.5 (61). We used *process_radtags* in Stacks 1.46 (62) on both the *Ta. granulosa* and *Th. sirtalis* datasets to demultiplex reads and removed sequences with low-quality scores or uncalled bases. For the *Ta. granulosa* dataset, we used the *denovo_map.pl* pipeline in Stacks, because a reference genome is not currently available. We used a minimum depth of three (-*m*), a distance of three between stacks (-*M*), and a distance of three between catalog loci (-*n*). The *Th. sirtalis* reads were aligned to the *Th. sirtalis* genome using Bowtie2 2.2.9 (63). We discarded reads that did not align or had more than one match to the genome. We used *ref_map.pl* in Stacks to assemble the reference-aligned sequences into loci, with a minimum depth of three (-*m*). For both species, we used *populations* in Stacks to select loci with a minimum depth of 10x coverage. To avoid linkage among sites within the same locus, we only retained one single nucleotide polymorphism (SNP) per locus.

We used the dartR package in R (64, 65) to remove loci and individuals with >30% missing data. We also removed loci with a minor allele frequency (MAF) <5%, including those that were invariant. Finally, we removed putative loci under selection. The program BayeScan 2.1 was used to search for loci with F_ST_ coefficients that were significantly different than those expected under neutrality (66). The Bayesian analysis used 20 pilot runs with 5,000 iterations followed by an additional burn-in of 50,000 and 50,000 output iterations. An outlier analysis with FDR-corrected p-values (q-values) <0.05 was used to identify and remove outlier loci putatively under selection.

Illumina HiSeq sequencing yielded an average of 1,093,529 raw reads per sample of *Ta. granulosa* (n=137). After initial quality control in *process_radtags*, the *denovo_map* pipeline in Stacks identified an average of 32,469 loci per individual, with a mean of 19x coverage. After further filtering steps in *populations*, dartR, and Bayescan, we retained 3,634 unlinked neutral SNPs in 123 individuals. For *Th. sirtalis*, sequencing yielded an average of 2,451,623 raw reads per sample (n=143). The *ref_map* pipeline identified an average of 13,501 loci per individual, with a mean of 80x coverage. After additional filtering, we retained 1,027 unlinked neutral SNPs in 132 individuals. The final datasets comprised a larger number of SNPs for *Ta. granulosa* (3, 634) than *Th. sirtalis* (1, 027), so we reran analyses with a random subsample of 1,027 SNPs for *Ta. granulosa* to generate equivalent datasets for each species. Results from the subsample were highly similar to the full dataset, so only results from the full analysis are presented herein.

### Analysis of geographic population structure

The filtered SNPs for each species were used to calculate observed heterozygosity (H_O_) and gene diversity (H_S_) for each population (67) using the hierfstat package in R (68). To estimate neutral population genomic structure, we calculated global and pairwise F_ST_ values (69). Confidence intervals were estimated by running 1000 bootstraps over loci using the hierfstat and stAMPP packages in R (70). Estimates of population genetic diversity from the final SNP dataset of each species are reported in Supplementary Table S3.

We tested for a pattern of isolation-by-distance (IBD) along each transect by performing Mantel tests on matrices of linearized pairwise F_ST_ and geographic distance (71). We also conducted distance-based redundancy analyses (dbRDA), which are thought to be more reliable than Mantel tests at detecting spatial patterns like IBD (72, 73). We conducted dbRDA analyses in the vegan package in R (74) to test for a relationship between pairwise F_ST_ values and the geographic coordinates (latitude and longitude) of sampling locations. We assessed significance of IBD tests with 1000 permutations.

We visualized population structure of each species using a principal coordinate analysis (PCoA) in the dartR package in R. The first axis (PCo1) from the PCoA captured latitudinal variation along the transect for both newts and snake (Fig. 3); therefore, we used the PCo1 values for each individual as a neutral expectation in the cline-fitting analyses (see below). We assessed population structure using a Bayesian assignment approach implemented in the program STRUCTURE (75, 76). We estimated the optimal number of genetic clusters (K) ranging for one to nine, without populations included as priors. The model assumed population admixture and correlated allele frequencies (76). The analysis first ran for 100,000 iterations as burn-in and then we collected data from the following 1,000,000 interactions in 10 different independent runs. STRUCTURE HARVESTER (77) was used to detect the most probable K using the Evanno’s method (78). Ancestry proportion (Q) values of the 10 runs for each value of the most probable K we averaged using CLUMPP (79) and visualized using the pophelper package in R (80). We calculated the average Q value for each population, which represent the fraction of membership to each genetic cluster (K). These Q estimates were used as a neutral expectation in cline-fitting analyses (see below) and compared to the cline results of PCo1 from the PCoA.

### Multiple regression of distance matrices (MRMs)

MRMs are an extension of Mantel tests that involve multiple regression of a response distance matrix on explanatory matrices (81–84). Partial regression coefficients can be used to understand the relationship between two matrices while controlling for the effects of a third matrix (7, 85). For each species, we generated two distance matrices: (1) pairwise phenotypic divergence in the coevolutionary trait (e.g., log-transformed TTX of *Ta. granulosa*) and (2) pairwise genomic divergence (F_ST_) from neutral SNPs. We then tested whether population patterns of phenotypic escalation in a focal species (e.g., TTX of *Ta. granulosa*) are best explained by neutral genomic divergence (pairwise F_ST_) or escalation in the natural enemy (TTX resistance of *Th. sirtalis*). MRM analyses can be confounded when explanatory variables are spatially autocorrelated, so we compared results when the two explanatory variables were analyzed separately and together (85–87).

### Cline analyses

We performed ML cline-fitting analyses on the phenotypic and genetic data from each species (88, 89). For *Ta. granulosa*, we fit a cline to TTX levels using the mean and variance of the log-transformed TTX data (μg/cm^2^). For *Th. sirtalis*, we fit (1) a cline to phenotypic TTX resistance using the 50% MAMU dose and variance from the ln-transformed MAMU data and (2) a genetic cline to the frequency of TTX-resistant alleles in each population. Two different TTX-resistant DIV alleles occur in the Pacific Northwest: Na_V_1.4^V^ is generally found at high frequency in the center of the transect, whereas Na_V_1.4^VA^ occurs predominately in southern populations (Fig. 1). We generated separate clines for each allele (data not shown), but the cline fit to the combined frequency of both TTX-resistant alleles (Fig. 4) was the most representative of variation along the entire transect.

For each species, we compared clinal variation in coevolutionary traits to variation at the neutral SNPs. We first fit clines to the PCo1 values from the PCoA of each species, and secondly, we fit clines to the ancestry proportions (Q) from the most likely value of K for each species (K=2). Both analyses of population structure (PCo1 from the PCoA and K=2 from STRUCTURE) produced concordant results (Supplemental Fig. S2; Supplemental Table S5), so only the analysis of PCo1 is presented in the main text. PCoAs have no explicit population genetic assumptions, whereas STRUCTURE assumes that all loci are unlinked at linkage equilibrium and in HWE. Because the PCoA and STRUCTURE analyses produced similar results despite different underlying assumptions, we feel confident that the cline-fitting analyses provided a representative depiction of neutral population structure in each species. Finally, we reran cline-fitting analyses with the Elk River population removed from the *Ta. granulosa* dataset, because the PCoA (Fig. 3) and STRUCTURE plot of K=3 (Supplemental Fig. S1) both suggest the population is genetically distinct from all others; however, this produced the same qualitative result as when the population was included.

We fit clines using the HZAR package in R (90). We calculated distances along the cline as kilometers (km) from the northernmost sampling site (Clallam). We ran 15 separate models that varied in the number of cline shape parameters estimated. All models estimated the cline center (distance from sampling location 1, *c*) and width (1/maximum slope, *w*), but could additionally estimate combinations of exponential decay curve (tail) parameters (neither tail, right tail only, left tail only, mirrored tails, or both tails separately), which represent the distance from the cline center to the tail (δ) and the slope of the tail (τ). The genetic models varied as to whether they estimated allele frequencies at the cline ends (*p*_min_ and *p*_max_) or fixed them at 0 and 1. All models were then compared using AIC corrected for small sample sizes (AICc) and maximum likelihood parameters were extracted for the best-fitting model. We considered cline centers with non-overlapping two log-likelihood unit support limits (confidence intervals; CIs) to occur in significantly different geographic locations (91, 92).

The cline-fitting analyses revealed a tight correlation between the TTX levels of *Ta. granulosa* and PCo1 from the PCoA of neutral SNPs (Fig. 4). To further assess this relationship, we fit linear models that tested whether clinal variation in TTX is best explained by neutral population structure from PCo1 and 2 of the PCoA (model 1) or escalation of TTX resistance in *Th. sirtalis* (model 2). We then fit a combined model including all explanatory variables (model 3) to test whether neutral structure from the PCoA predicts TTX levels even after controlling for the TTX resistance of *Th. sirtalis* (Supplemental Table S6).

We conducted an analogous set of tests (models 1-3) for *Th. sirtalis* to assess whether TTX resistance is best predicted by neutral structure from the PCoA or the TTX levels of *Ta. granulosa*; however, the analyses were more complicated because population estimates of TTX resistance (50% MAMU doses) are extracted from a dose response curve. TTX resistance of individual snakes is measured as a proportion (i.e., the reduction in crawl speed after a given dose of TTX), with each individual receiving multiple injections. We accounted for these data by fitting linear mixed models using the “lmer” function in the R package *lme4* (47). We treated “post-injection” crawl speed (after a given dose of TTX) as the response variable, and included “baseline” crawl speed and the dose of TTX as covariates in each model. We also included individual ID as a random effect to account for the fact that individuals received multiple doses. Post-injection crawl speed was log-transformed to account for non-normality. Statistical significance of fixed effects was determined by an ANOVA using a Wald Chi-Square test with type III sum of squares and one degree of freedom, implemented in the *car* package in R (93). We reported the marginal R^2^ for each model, which describes the proportion of variance explained by the fixed factors (not including random effects), implemented in the *MuMIn* package (94, 95).

### Data availability

DNA sequence alignments for the DIV p-loop of Na_V_1.4 and the ddRADseq data will be made available on GenBank upon manuscript acceptance. All phenotypic data and the code for statistical analyses will be submitted to Dryad upon acceptance.

## SUPPLEMENTAL INFORMATION

**Supplemental Fig. S1.** STRUCTURE results for K=2-4. The most likely number of genetic cluster was K=2 for both species (Fig. 2). STRUCTURE plots are arranged latitudinally by population, in the same order as the map. Each horizontal bar represents the ancestry assignment of an individual, with populations separated by white dashed lines.

**Supplemental Fig. S2.** Comparison of neutral clines fit to (1) PCo1 values from the PCoA and (2) average ancestry assignment values (K=2) from the STRUCTURE analysis. Error bars indicate confidence intervals surrounding the geographic cline centers.

---

**Supplemental Table S1.** Datasets for each sampling location along the latitudinal transect. The total number of animals sampled in this study is shown first, followed by the number of individuals (*n*) included in each analysis. For estimates of phenotypic resistance in *Th. sirtalis*, we combined individuals from this study with previously published racetrack data from the same sampling locations (14, 45). In the DIV p-loop, the number (*n*) of alleles sampled accounts for females, the hemizygous sex with only one genetic copy of Na_V_1.4. Results are shown for the joint test of Hardy-Weinberg Equilibrium (HWE) and Equality of Allele Frequencies (EAF).

| Location | County | Total animals sampled<br>in this study |  | *Ta. granulosa* |  |  |  | *Th. sirtalis* |  |  |  |  |  |
| --- | --- | --- | --- | --- | --- | --- | --- | --- | --- | --- | --- | --- | --- |
|  |  |  |  | TTX |  |  | ddRADseq | Phenotypic TTX resistance |  |  | DIV p-loop |  | ddRADseq |
|  |  | *Ta.*<br>*granulosa* | *Th.*<br>*sirtalis* | *n* | mean<br>(ug/cm<sup>2</sup>) | SE | *n* | *n* | 50%<br>MAMU | SE | *n*<br>alleles | HWE &<br>EAF test<br>p-value | *n* |
| Clallam | Clallam, WA | 12 | 11 | 12 | 0.731 | 0.307 | 12 | 53 | 4.163 | 0.076 | 14 | NA | 4 |
| Cook Creek | Grays Harbor, WA | 14 | 17 | 14 | 0.236 | 0.112 | 13 | 14 | 5.915 | 0.119 | 26 | 1.000 | 16 |
| Potter's Slough | Pacific, WA | 15 | 20 | 15 | 2.75 | 0.686 | 13 | 42 | 19.203 | 0.090 | 27 | 1.000 | 16 |
| Warrenton | Clatsop, OR | 15 | 24 | 14 | 1.63 | 0.241 | 14 | 323 | 17.357 | 0.038 | 36 | 0.257 | 20 |
| Hebo | Tillamook, OR | 15 | 15 | 15 | 4.65 | 0.822 | 12 | 2 | 43.413 | 0.085 | 21 | 0.708 | 14 |
| Benton | Benton., OR | 18 | 20 | 18 | 16.9 | 3.1 | 17 | 386 | 31.380 | 0.047 | 23 | 0.347 | 14 |
| Ten Mile | Lane, OR | 15 | 16 | 14 | 7.43 | 1.85 | 14 | 93 | 52.817 | 0.079 | 26 | 0.451 | 16 |
| Lake<br>Tahkenitch | Douglas, OR | 14 | 29 | 13 | 10.7 | 2.56 | 12 | 17 | 14.552 | 0.227 | 42 | 0.182 | 15 |
| Elk River | Curry, OR | 20 | 17 | 15 | 6.52 | 0.612 | 16 | 43 | 12.935 | 0.063 | 20 | 0.324 | 17 |
| Total |  | 138 | 169 | 130 |  |  | 123 | 932 |  |  | 235 |  | 132 |

**Supplemental Table S2.** For each locality, functional estimates of adult *Ta. granulosa* total skin TTX (mg) and oral doses of TTX (mg) required to reduce the speed of an average adult *Th. sirtalis* to 15, 50, and 85% of baseline speed post-ingestion of TTX.

| Population | *Ta. granulosa* |  |  | *Th. sirtalis* |  |  |
| --- | --- | --- | --- | --- | --- | --- |
|  | mean TTX<br>(mg) | max TTX<br>(mg) | min TTX<br>(mg) | 50% oral<br>dose (mg) | 15% oral<br>dose (mg) | 85% oral<br>dose (mg) |
| Clallam | 0.0176 | 0.0715 | 0.0001 | 0.1257 | 0.5710 | 0.0102 |
| Cook Creek | 0.0068 | 0.0422 | 0.0001 | 0.1787 | 0.4614 | 0.0585 |
| Potter's Slough | 0.0816 | 0.3813 | 0.0220 | 0.5800 | 1.9169 | 0.1610 |
| Warrenton | 0.0475 | 0.1263 | 0.0053 | 0.5243 | 2.2042 | 0.1074 |
| Hebo | 0.1933 | 0.5495 | 0.0698 | 1.3113 | 1.7279 | 0.9935 |
| Benton | 0.7107 | 1.9188 | 0.1097 | 0.9479 | 3.9647 | 0.2093 |
| Ten Mile | 0.3188 | 1.0166 | 0.0455 | 1.5954 | 11.1998 | 0.2051 |
| Lake Tahkenitch | 0.2590 | 0.8172 | 0.0468 | 0.4396 | 3.4492 | 0.0332 |
| Elk River | 0.2043 | 0.3495 | 0.1007 | 0.3907 | 1.4213 | 0.0918 |

**Supplemental Table S3.** Population genetic diversity statistics from neutral SNPs in each species. Sample size (*n*), average observed heterozygosity (H_O_), and expected heterozygosity (H_E_) are shown, along with standard deviations (SD).

| Population | *Ta. granulosa* |  |  |  |  | *Th. sirtalis* |  |  |  |  |
| --- | --- | --- | --- | --- | --- | --- | --- | --- | --- | --- |
|  | *n* | *H<sub>O</sub>* | *H<sub>O</sub>* SD | *H<sub>E</sub>* | *H<sub>S</sub>* SD | *n* | *H<sub>O</sub>* | *H<sub>O</sub>* SD | *H<sub>E</sub>* | *H<sub>S</sub>* SD |
| Clallam | 12 | 0.278 | 0.248 | 0.253 | 0.196 | 4 | 0.328 | 0.297 | 0.294 | 0.218 |
| Cook Creek | 13 | 0.324 | 0.237 | 0.294 | 0.178 | 16 | 0.311 | 0.231 | 0.288 | 0.177 |
| Potters Slough | 13 | 0.352 | 0.225 | 0.320 | 0.163 | 16 | 0.319 | 0.230 | 0.293 | 0.169 |
| Warrenton | 14 | 0.321 | 0.200 | 0.323 | 0.156 | 20 | 0.297 | 0.218 | 0.292 | 0.170 |
| Hebo | 12 | 0.330 | 0.205 | 0.332 | 0.159 | 14 | 0.292 | 0.218 | 0.302 | 0.168 |
| Benton | 17 | 0.335 | 0.198 | 0.330 | 0.154 | 14 | 0.337 | 0.270 | 0.288 | 0.184 |
| Ten Mile | 14 | 0.325 | 0.201 | 0.332 | 0.157 | 16 | 0.363 | 0.246 | 0.303 | 0.163 |
| Tahkenitch | 12 | 0.376 | 0.244 | 0.332 | 0.169 | 15 | 0.303 | 0.220 | 0.296 | 0.169 |
| Elk River | 16 | 0.300 | 0.204 | 0.318 | 0.174 | 17 | 0.306 | 0.232 | 0.285 | 0.180 |
| Total | 123 | 0.327 | 0.218 | 0.315 | 0.168 | 132 | 0.317 | 0.240 | 0.293 | 0.178 |

**Supplemental Table S4.** Pairwise F_ST_ statistics for the neutral SNP datasets of each species. For *Th. sirtalis*, F_ST_ divergence at the DIV p-loop of the Na_V_1.4 channel is also shown. F_ST_ values are shaded red to illustrate the extent of divergence among different populations (white=0.00, red=1.00).

| *Ta. granulosa* pairwise $F_{ST}$ of neutral SNPs | | | | | | | | |
| --- | --- | --- | --- | --- | --- | --- | --- | --- |
|  | Clallam | Cook Creek | Potters Slough | Warrenton | Hebo | Benton | Ten Mile | Tahkenitch |
| Cook Creek | 0.040 |  |  |  |  |  |  |  |
| Potters Slough | 0.066 | 0.011 |  |  |  |  |  |  |
| Warrenton | 0.077 | 0.024 | 0.004 |  |  |  |  |  |
| Hebo | 0.111 | 0.058 | 0.031 | 0.019 |  |  |  |  |
| Benton | 0.158 | 0.096 | 0.057 | 0.039 | 0.020 |  |  |  |
| Ten Mile | 0.147 | 0.088 | 0.053 | 0.034 | 0.013 | 0.007 |  |  |
| Tahkenitch | 0.144 | 0.088 | 0.055 | 0.040 | 0.022 | 0.016 | 0.009 |  |
| Elk River | 0.218 | 0.154 | 0.116 | 0.101 | 0.081 | 0.066 | 0.065 | 0.067 |
| *Th. sirtalis* pairwise $F_{ST}$ of neutral SNPs | | | | | | | | |
|  | Clallam | Cook Creek | Potters Slough | Warrenton | Hebo | Benton | Ten Mile | Tahkenitch |
| Cook Creek | 0.044 |  |  |  |  |  |  |  |
| Potters Slough | 0.058 | 0.025 |  |  |  |  |  |  |
| Warrenton | 0.069 | 0.054 | 0.044 |  |  |  |  |  |
| Hebo | 0.067 | 0.063 | 0.051 | 0.052 |  |  |  |  |
| Benton | 0.120 | 0.104 | 0.084 | 0.082 | 0.045 |  |  |  |
| Ten Mile | 0.066 | 0.070 | 0.058 | 0.062 | 0.024 | 0.056 |  |  |
| Tahkenitch | 0.097 | 0.087 | 0.075 | 0.086 | 0.031 | 0.065 | 0.029 |  |
| Elk River | 0.150 | 0.129 | 0.114 | 0.127 | 0.083 | 0.110 | 0.064 | 0.031 |
| *Th. sirtalis* pairwise $F_{ST}$ of DIV p-loop | | | | | | | | |
|  | Clallam | Cook Creek | Potters Slough | Warrenton | Hebo | Benton | Ten Mile | Tahkenitch |
| Cook Creek | 0.039 |  |  |  |  |  |  |  |
| Potters Slough | 0.688 | 0.625 |  |  |  |  |  |  |
| Warrenton | 0.722 | 0.665 | 0.014 |  |  |  |  |  |
| Hebo | 0.575 | 0.497 | 0.034 | 0.154 |  |  |  |  |
| Benton | 0.652 | 0.579 | 0.025 | 0.162 | -0.034 |  |  |  |
| Ten Mile | 0.593 | 0.514 | 0.105 | 0.251 | -0.028 | -0.019 |  |  |
| Tahkenitch | 0.820 | 0.739 | 0.455 | 0.595 | 0.254 | 0.245 | 0.147 |  |
| Elk River | 0.805 | 0.725 | -0.010 | -0.024 | 0.141 | 0.132 | 0.230 | 0.603 |

**Supplemental Table S5.** Results from cline-fitting analyses. The mean value and its geographic center point along the cline are shown for each dataset, in addition to cline width. Confidence intervals (CI) are also listed.

|  |  | Mean | Center (km) | Lower CI | Upper CI | Width (km) | Lower CI | Upper CI |
| --- | --- | --- | --- | --- | --- | --- | --- | --- |
| *Ta. granulosa* | **TTX** | 0.118 | 193.184 | 168.778 | 200.283 | 284.949 | 207.732 | 363.750 |
|  | **Neutral PCo1** | -2.562 | 212.131 | 158.176 | 250.096 | 599.042 | 532.016 | 611.000 |
|  | **STRUCTURE (K=2)** | 0.500 | 229.646 | 188.078 | 265.566 | 440.284 | 348.296 | 574.741 |
| *Th. sirtalis* | **50% MAMU dose** | 2.350 | 128.543 | 113.922 | 173.242 | 510.820 | 314.218 | 539.450 |
|  | **Freq. of TTX resistant alleles** | 0.500 | 128.843 | 104.118 | 149.925 | 55.589 | 5.569 | 101.699 |
|  | **Neutral PCo1** | 1.120 | 439.140 | 412.333 | 501.383 | 553.836 | 411.980 | 610.930 |
|  | **STRUCTURE (K=2)** | 0.506 | 387.698 | 359.552 | 415.000 | 276.425 | 206.047 | 368.235 |

**Supplemental Table S6.**
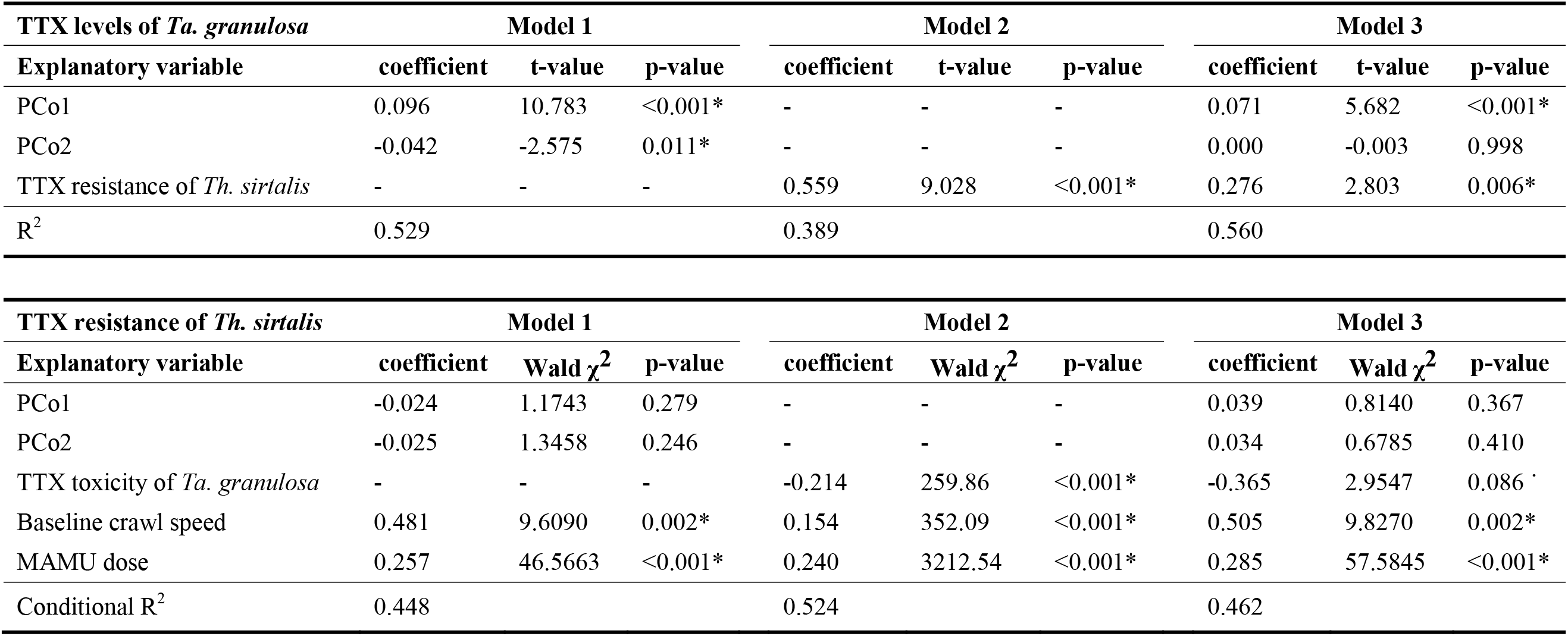
Results from the linear models (*Ta. granulosa*) and linear mixed models (*Th. sirtalis*) accompanying the cline analyses. Separate models were fit to test whether variation in each coevolutionary trait is predicted by neutral population structure from the PCoA (model 1) or phenotypic variation in the natural enemy (model 2). Model 3 includes all explanatory variables. The TTX resistance of *Th. sirtalis* was measured as post-injection crawl speed, with baseline crawl speed and MAMU dose included in the model as covariates.

